# A model species for agricultural pest genomics: the genome of the Colorado potato beetle, *Leptinotarsa decemlineata* (Coleoptera: Chrysomelidae)

**DOI:** 10.1101/192641

**Authors:** Sean D. Schoville, Yolanda H. Chen, Martin N. Andersson, Joshua B. Benoit, Anita Bhandari, Julia H. Bowsher, Kristian Brevik, Kaat Cappelle, Mei-Ju M. Chen, Anna K. Childers, Christopher Childers, Olivier Christiaens, Justin Clements, Elise M. Didion, Elena N. Elpidina, Patamarerk Engsontia, Markus Friedrich, Inmaculada García-Robles, Richard A. Gibbs, Chandan Goswami, Alessandro Grapputo, Kristina Gruden, Marcin Grynberg, Bernard Henrissat, Emily C. Jennings, Jeffery W. Jones, Megha Kalsi, Sher A. Khan, Abhishek Kumar, Fei Li, Vincent Lombard, Xingzhou Ma, Alexander Martynov, Nicholas J. Miller, Robert F. Mitchell, Monica Munoz-Torres, Anna Muszewska, Brenda Oppert, Subba Reddy Palli, Kristen A. Panfilio, Yannick Pauchet, Lindsey C. Perkin, Marko Petek, Monica F. Poelchau, Éric Record, Joseph P. Rinehart, Hugh M. Robertson, Andrew J. Rosendale, Victor M. Ruiz-Arroyo, Guy Smagghe, Zsofia Szendrei, Gregg W. C. Thomas, Alex S. Torson, Iris M. Vargas Jentzsch, Matthew T. Weirauch, Ashley D. Yates, George D. Yocum, June-Sun Yoon, Stephen Richards

**Affiliations:** Department of Entomology, University of Wisconsin-Madison; Department of Plant and Soil Sciences, University of Vermont; Department of Biology, Lund University; Department of Biological Sciences, University of Cincinnati; Department of Molecular Physiology, Christian-Albrechts-University at Kiel; Department of Biological Sciences, North Dakota State University; Department of Crop Protection, Ghent University; USDA-ARS National Agricultural Library, Beltsville, MD USA; USDA-ARS Bee Research Lab, Beltsville, MD USA; USDA-ARS Insect Genetics and Biochemistry Research Unit, Fargo, ND USA; A.N. Belozersky Institute of Physico-Chemical Biology, Lomonosov Moscow State University; Department of Biology, Faculty of Science, Prince of Songkla University, Thailand; Department of Biological Sciences, Wayne State University; Department of Genetics, University of Valencia; Department of Molecular and Human Genetics, Baylor College of Medicine; National Institute of Science Education and Research, Bhubaneswar, India; Department of Biotechnology and Systems Biology, National Institute of Biology, Slovenia; Institute of Biochemistry and Biophysics, Polish Academy of Sciences; Architecture et Fonction des Macromolécules Biologiques, CNRS, Aix-Marseille Université, 13288 Marseille, France; INRA, USC 1408 AFMB, F-13288 Marseille, France; Department of Biological Sciences, King Abdulaziz University, Saudi Arabia; Department of Entomology, University of Kentucky; Department of Entomology, Texas A&M University, College Station; Department of Genetics & Molecular Biology in Botany, Christian-Albrechts-University at Kiel; Division of Molecular Genetic Epidemiology, German Cancer Research Center, Heidelberg; Department of Entomology, Nanjing Agricultural University; Center for Data-Intensive Biomedicine and Biotechnology, Skolkovo Institute of Science and Technology; Department of Biology, Illinois Institute of Technology; Department of Biology, University of Wisconsin-Oshkosh; Environmental Genomics and Systems Biology Division, Lawrence Berkeley National Laboratory; USDA-ARS Center for Grain and Animal Health Research; Institute for Developmental Biology, University of Cologne; School of Life Sciences, University of Warwick, Gibbet Hill Campus; Department of Entomology, Max Planck Institute for Chemical Ecology, Jena, Germany; INRA, Aix-Marseille Université, UMR1163, Biodiversité et Biotechnologie Fongiques, Marseille, France; Department of Entomology, University of Illinois at Urbana-Champaign; Department of Entomology, Michigan State University; Department of Biology and School of Informatics and Computing, Indiana University; Center for Autoimmune Genomics and Etiology, Division of Biomedical Informatics and Division of Developmental Biology, Cincinnati Children’s Hospital Medical Center.; Department of Pediatrics, University of Cincinnati College of Medicine.; Department of Entomology, The Ohio State University; Center for Applied Plant Sciences, The Ohio State University

**Keywords:** insects, evolution, pesticide resistance, herbivory, plant-insect interactions, pest management, whole-genome sequence

## Abstract

The Colorado potato beetle is one of the most challenging agricultural pests to manage. It has shown a spectacular ability to adapt to a variety of solanaceaeous plants and variable climates during its global invasion, and, notably, to rapidly evolve insecticide resistance. To examine evidence of rapid evolutionary change, and to understand the genetic basis of herbivory and insecticide resistance, we tested for structural and functional genomic changes relative to other arthropod species using genome sequencing, transcriptomics, and community annotation. Two factors that might facilitate rapid evolutionary change include transposable elements, which comprise at least 17% of the genome and are rapidly evolving compared to other Coleoptera, and high levels of nucleotide diversity in rapidly growing pest populations. Adaptations to plant feeding are evident in gene expansions and differential expression of digestive enzymes in gut tissues, as well as expansions of gustatory receptors for bitter tasting. Surprisingly, the suite of genes involved in insecticide resistance is similar to other beetles. Finally, duplications in the RNAi pathway might explain why *Leptinotarsa decemlineata* has high sensitivity to dsRNA. The *L. decemlineata* genome provides opportunities to investigate a broad range of phenotypes and to develop sustainable methods to control this widely successful pest.

## Introduction

The Colorado potato beetle, *Leptinotarsa decemlineata* Say 1824 (Coleoptera: Chrysomelidae), is widely considered one of the world’s most successful globally-invasive insect herbivores, with costs of ongoing management reaching tens of millions of dollars annually [1] and projected costs if unmanaged reaching billions of dollars [2]. This beetle was first identified as a pest in 1859 in the Midwestern United States, after it expanded from its native host plant, *Solanum rostratum* (Solanaceae), onto potato (*S. tuberosum*) [3]. As testimony to the difficulty in controlling *L. decemlineata*, the species has the dubious honor of starting the pesticide industry, when Paris Green (copper (II) acetoarsenite) was first applied to control it in the United States in 1864 [4]. *Leptinotarsa decemlineata* is now widely recognized for its ability to rapidly evolve resistance to insecticides, as well as a wide range of abiotic and biotic stresses [5], and for its global expansion across 16 million km^2^ to cover the entire Northern Hemisphere within the 20th century [6]. Over the course of 150 years of research, *L. decemlineata* has been the subject in more than 9,700 publications (according to the Web of Science™ Core Collection of databases) ranging from molecular to organismal biology from the fields of agriculture, entomology, molecular biology, ecology, and evolution.

In order to be successful, *L. decemlineata* evolved to exploit novel host plants, to inhabit colder climates at higher latitudes [7–9], and to cope with a wide range of novel environmental conditions in agricultural landscapes [10,11]. Genetic data suggest the potato-feeding pest lineage directly descended from populations that feed on *S. rostratum* in the U.S. Great Plains [12]. This beetle subsequently expanded its range northwards, shifting its life history strategies to exploit even colder climates [7,8,13], and steadily colonized potato crops despite substantial geographical barriers [14]. *Leptinotarsa decemlineata* is an excellent model for understanding pest evolution in agroecosystems because, despite its global spread, individuals disperse over short distances and populations often exhibit strong genetic differentiation [15–17], providing an opportunity to track the spread of populations and the emergence of novel phenotypes. The development of genomic resources in *L. decemlineata* will provide an unparalleled opportunity to investigate the molecular basis of traits such as climate adaptation, herbivory, host expansion, and chemical detoxification. Perhaps most significantly, understanding its ability to evolve rapidly would be a major step towards developing sustainable methods to control this widely successful pest in agricultural settings.

Given that climate is thought to be the major factor in structuring the range limits of species [18], the latitudinal expansion of *L. decemlineata*, spanning more than 40° latitude from Mexico to northern potato-producing countries such as Canada and Russia [6], warrants further investigation. Harsh winter climates are thought to present a major barrier for insect range expansions, especially near the limits of a species’ range [7,19]. To successfully overwinter in temperate climates, beetles need to build up body mass, develop greater amounts of lipid storage, have a low resting metabolism, and respond to photoperiodic keys by initiating diapause [20,21]. Although the beetle has been in Europe for less than 100 years, local populations have demonstrating remarkably rapid evolution in life history traits linked to growth, diapause and metabolism [8,13,20]. Understanding the genetic basis of these traits, particularly the role of specific genes associated with metabolism, fatty acid synthesis, and diapause induction, could provide important information about the mechanism of climate adaptation.

Although *Leptinotarsa decemlineata* has long-served as a model for the study of host expansion and herbivory due to its rapid ability to host switch [17,22], a major outstanding question is what genes and biological pathways are associated with herbivory in this species?

While >35,000 species of Chrysomelidae are well-known herbivores, most species feed on one or a few host species within the same plant family [23]. Within *Leptinotarsa*, the majority of species feed on plants within Solanaceae and Asteraceae, while *L. decemlineata* feeds exclusively on solanaceous species [24]. It has achieved the broadest host range amongst its congeners (including, but not limited to: buffalobur (*S. rostratum*), potato (*S. tuberosum*), eggplant (*S. melongena*), silverleaf nightshade (*S. elaeagnifolium*), horsenettle (*S. carolinense*), bittersweet nightshade (S. *dulcamara*), tomato (*S. lycopersicum*), and tobacco (*Nicotiana tabacum*)) [17,22,25], and exhibits geographical variation in the use of locally abundant *Solanum* species [26].

Another major question is what are the genes that underlie the beetle’s remarkable capacity to detoxify plant secondary compounds and are these the same biological pathways used to detoxify insecticidal compounds [27]? Solanaceous plants are considered highly toxic to a wide range of insect herbivore species [28], because they contain steroidal alkaloids and glycoalkaloids, nitrogen-containing compounds that are toxic to a wide range of organisms, including bacteria, fungi, humans, and insects [29], as well as glandular trichomes that contain additional toxic compounds [30]. In response to beetle feeding, potato plants upregulate pathways associated with terpenoid, alkaloid, and phenylpropanoid biosynthesis, as well as a range of protease inhibitors [31]. A complex of digestive cysteine proteases is known to underlie *L. decemlineata*’s ability to respond to potato-induced defenses [32,33]. There is evidence that larvae excrete [34] and perhaps even sequester toxic plant-based compounds in the hemolymph [35,36]. Physiological mechanisms involved in detoxifying plant compounds, as well as other xenobiotics, have been proposed to underlie pesticide resistance [27]. To date, while cornerstone of *L. decemlineata* management has been the use of insecticides, the beetle has evolved resistance to over 50 compounds and all of the major classes of insecticides. Some of these chemicals have even failed to control *L. decemlineata* within the first year of release [10], and notably, regional populations of *L. decemlineata* have demonstrated the ability to independently evolve resistance to pesticides and to do so at different rates [37]. Previous studies have identified target site mutations in resistance phenotypes and a wide range of genes involved in metabolic detoxification, including carboxylesterase genes, cytochrome P450s, and glutathione S-transferase genes [38–42].

To examine evidence of rapid evolutionary change underlying *L. decemlineata’s* extraordinary success utilizing novel host plants, climates, and detoxifying insecticides, we evaluated structural and functional genomic changes relative to other beetle species, using whole-genome sequencing, transcriptome sequencing, and a large community-driven biocuration effort to improve predicted gene annotations. We compared the size of gene families associated with particular traits against existing available genomes from related species, particularly those sequenced by the i5k project (http://i5k.github.io), an initiative to sequence 5,000 species of Arthropods. While efforts have been made to understand the genetic basis of phenotypes in *L. decemlineata* (for example, pesticide resistance) [32,43,44], previous work has been limited to candidate gene approaches rather than comparative genomics. Genomic data can not only illuminate the genetic architecture of a number of phenotypic traits that enable *L. decemlineata* to continue to be an agricultural pest, but can also be used to identify new gene targets for control measures. For example, recent efforts have been made to develop RNAi-based pesticides targeting critical metabolic pathways in *L. decemlineata* [41,45,46]. With the extensive wealth of biological knowledge and a newly-released genome, this beetle is well-positioned to be a model system for agricultural pest genomics and the study of rapid evolution.

## Results and Discussion

### Genome Assembly, Annotation and Assessment

A single female *L. decemlineata* from Long Island, NY, USA, a population known to be resistant to a wide range of insecticides [47,48], was sequenced to a depth of ~140x coverage and assembled with ALLPATHS [49] followed by assembly improvement with ATLAS (https://www.hgsc.bcm.edu/software/). The average coleopteran genome size is 760 Mb (ranging from 160-5,020 Mb [50]), while most of the beetle genome assemblies have been smaller (mean assembly size 286 Mb, range 160-710 Mb) [51–55]. The draft genome assembly of *L. decemlineata* is 1.17 Gb and consists of 24,393 scaffolds, with a N50 of 414 kb and a contig N50 of 4.9 kb. This assembly is more than twice the estimated genome size of 460 Mb [56], with the presence of gaps comprising 492 Mb, or 42%, of the assembly. As this size might be driven by underlying heterozygosity, we also performed scaffolding with REDUNDANS [57], which reduced the assembly size to 642 Mb, with gaps reduced to 1.3% of the assembly. However, the REDUNDANS assembly increased the contig N50 to 47.4 kb, the number of scaffolds increased to 90,205 and the N50 declined to 139 kb. By counting unique 19 bp kmers and adjusting for ploidy, we estimate the genome size as 816.9 Mb. Using just the small-insert (180 bp) 100 bp PE reads, average coverage was 24X for the ALLPATHS assembly and 26X for the REDUNDANS assembly. For all downstream analyses, the ALLPATHS assembly was used due to its increased scaffold length and reduced number of scaffolds.

The number of genes in the *L. decemlineata* genome predicted based on automated annotation using MAKER was 24,671 gene transcripts, with 93,782 predicted exons, which surpasses the 13,526-22,253 gene models reported in other beetle genome projects (**Figure 1**) [51–55]. This may be in part due to fragmentation of the genome, which is known to inflate gene number estimates [58]. To improve our gene models, we manually annotated genes using expert opinion and additional mRNA resources (see **Supplementary File** for more details). A total of 1,364 genes were manually curated and merged with the unedited MAKER annotations to produce an official gene set (OGS v1.0) of 24,850 transcripts, comprised of 94,859 exons. A total of 12 models were curated as pseudogenes. A total of 1,237 putative transcription factors (TFs) were identified in the *L. decemlineata* predicted proteome (**Figure 2**). The predicted number of TFs is similar to some beetles, such as *Anoplophora glabripennis* (1,397) and *Hypothenemus hampei* (1,148), but substantially greater than others, such as *Tribolium castaneum* (788), *Nicrophorus vespilloides* (744), and *Dendroctonus ponderosae* (683) [51–55].

**Figure 1.**
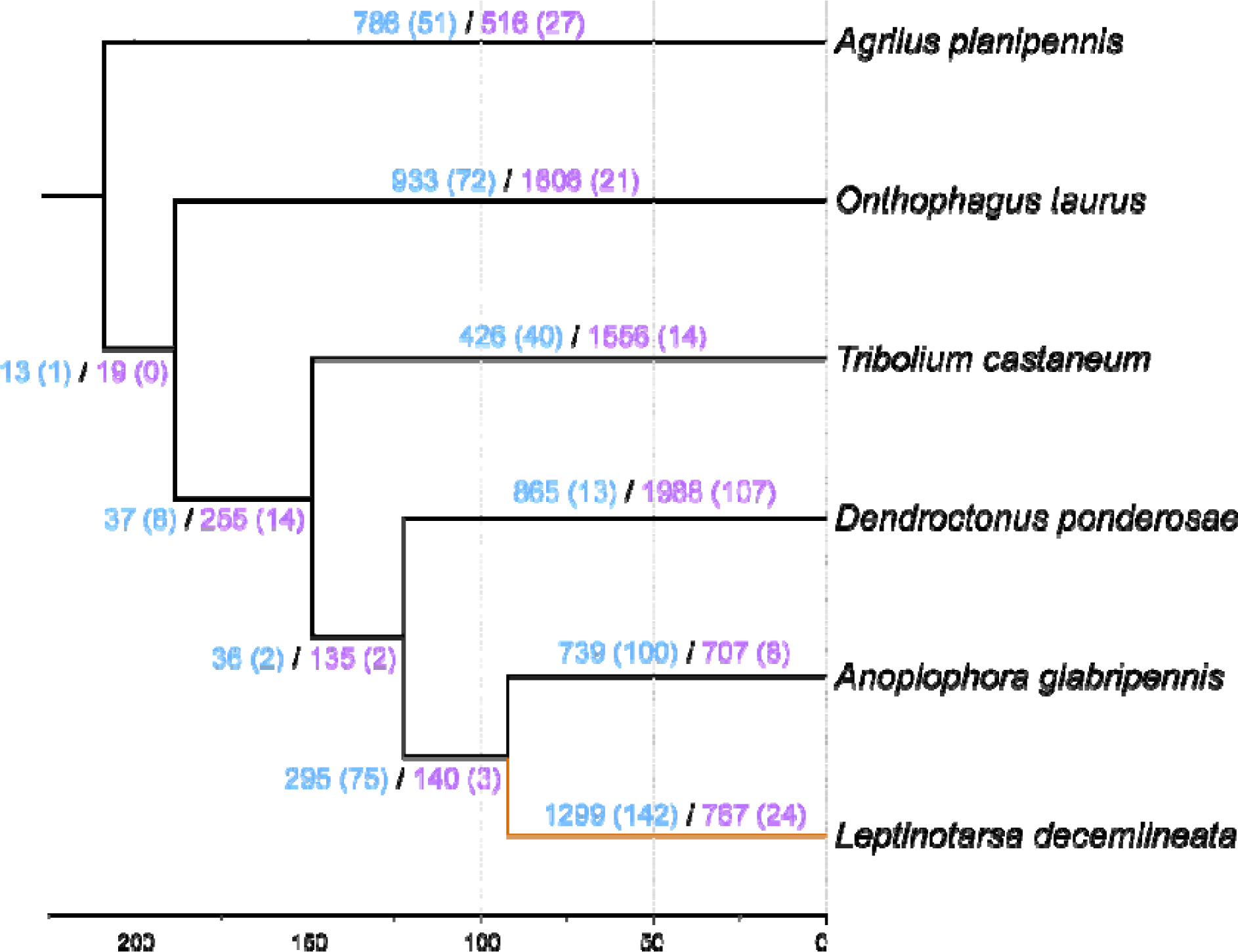
Ultrametric tree with branch lengths in millions of years for *Leptinotarsa decemlineata* relative to five other Coleoptera genomes. The *L. decemlineata* lineage is shown in orange. Branches are labeled with their length in years (top) and with the number of gene family expansions (blue) and contractions (purple) that occurred on that lineage. Rapid changes for both types are in parentheses.

**Figure 2.**
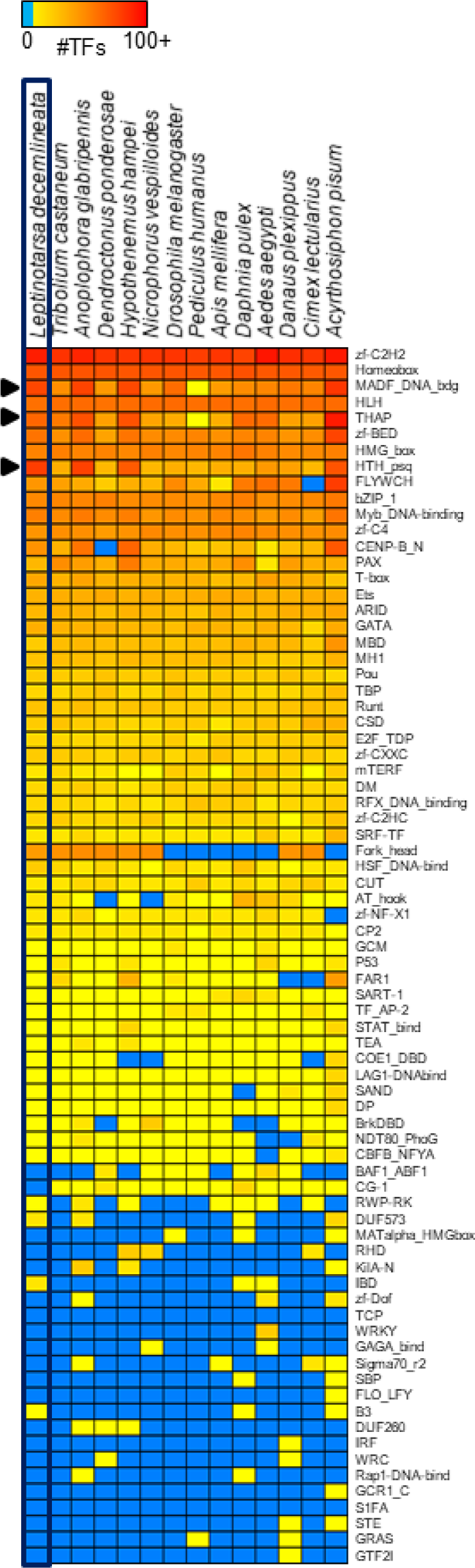
Heatmap distribution of the abundance of transcription factor families in *Leptinotarsa decemlineata* compared to other insects. Each entry indicates the number of TF genes for the given family per genome, based on presence of predicted DNA binding domains. Color scale is log (base 2) and the key is depicted at the top (light blue means the TF family is completely absent). Families discussed in the main text are indicated by arrows.

We assessed the completeness of both the ALLPATHS and REDUNDANS assemblies, and the OGS separately, using benchmarking sets of universal single-copy orthologs (BUSCOs) based on 35 holometabolous insect genomes [59], as well as manually assessing the completeness and co-localization of the homeodomain transcription factor gene clusters in the ALLPATHS assembly. Using the reference set of 2,442 BUSCOs, the ALLPATHS genome assembly, REDUNDANS genome assembly, and OGS were 93.0%, 91.9%, and 71.8% complete, respectively. We found an additional 4.1%, 5.4%, and 17.9% of the BUSCOs present but fragmented in each dataset, respectively. For the highly conserved Hox and Iroquois Complex (Iro-C) clusters, we located and annotated complete gene models in the ALLPATHS genome assembly for all 12 expected orthologs, but these were split across six different scaffolds (**Supplementary Figure 1S** and **Table 1S**). All linked Hox genes occurred in the expected order and with the expected, shared transcriptional orientation, suggesting that the current draft assembly was correct but incomplete (see also **Supplementary Figures 2S** and **3S**). Assuming direct concatenation of scaffolds, the Hox cluster would span a region of 3.7 Mb, similar to the estimated 3.5 Mb Hox cluster of *A. glabripennis* [54]. While otherwise highly conserved with *A. glabripennis*, we found a tandem duplication for *Hox3/zen* and an Antennapedia-class (ANTP-class) homeobox gene with no clear ortholog in other arthropods. We also assessed the ALLPATHS genome assembly for evidence of contamination using a Blobplot (**Supplementary Figure 4S**), which identified a small proportion of the reads (5.9%) as putative contaminants (the largest single taxonomic group represented, 2.6%, was assigned to Annelida).

### Gene Annotation, Gene Family Evolution and Differential Expression

We estimated a phylogeny among six coleopteran genomes (*A. glabripennis, Agrilus planipennis, D. ponderosae, L. decemlineata, Onthophagus taurus,* and *T. castaneum;* unpublished genomes available at http://i5k.github.io/) [51,52,54] using a conserved set of single copy orthologs and compared the official gene set of each species to understand how gene families evolved the branch representing Chysomelidae. *Leptinotarsa decemlineata* and *A. glabripennis* (Cerambycidae) are sister taxa (**Figure 1**), as expected for members of the same superfamily Chrysomeloidea. We found 166 rapidly evolving gene families along the *L. decemlineata* lineage (1.4% of 11,598), 142 of which are rapid expansions and the remaining 24 rapid contractions (**Table 1**). Among all branches of our coleopteran phylogeny, *L. decemlineata* has the highest average expansion rate (0.203 genes per million years), the highest number of genes gained, and the greatest number of rapidly evolving gene families.

**Table 1:** Summary of gene gain and loss events inferred after correcting for annotation and assembly error across six Coleoptera species. The number of rapidly evolving families is shown in parentheses for each type of change and the rate is genes per million years.

|  | Expansions |  |  | Contractions |  |  | No Change | Average Expansion |
| --- | --- | --- | --- | --- | --- | --- | --- | --- |
|  | Families | Genes gained | Rate | Families | Genes lost | Rate |  |  |
| *Anoplophora glabripennis* | 865 (13) | 1231 | 1.42 | 1988 (107) | 3125 | 1.57 | 5850 | -0.182341 |
| *Agrilus plannipennis* | 739 (100) | 1991 | 2.69 | 707 (8) | 769 | 1.09 | 7257 | 0.119108 |
| *Dendroctonus ponderosae* | 933 (72) | 1982 | 2.12 | 1606 (21) | 1887 | 1.17 | 6164 | 0.006438 |
| *Leptinotarsa decemlineata* | 426 (40) | 855 | 2.01 | 1556 (48) | 2116 | 1.36 | 6721 | -0.127501 |
| *Onthophagus taurus* | 1299 (142) | 2952 | 2.27 | 767 (24) | 895 | 1.17 | 6637 | 0.203380 |
| *Tribolium castaneum* | 786 (51) | 1428 | 1.82 | 516 (27) | 909 | 1.76 | 7401 | 0.055645 |

Examination of the functional classification of rapidly evolving families in *L. decemlineata* (**Supplementary Tables 2S** and **3S**) indicates that a subset of families are clearly associated with herbivory. The peptidases, comprising several gene families that play a major role in plant digestion [32,60], displayed a significant expansion in genes (OrthoDB family Id: EOG8JDKNM, EOG8GTNV8, EOG8DRCSN, EOG8GTNT0, EOG8K0T47, EOG88973C, EOG854CDR, EOG8Z91BB, EOG80306V, EOG8CZF0X, EOG80P6ND, EOG8BCH22, EOG8ZKN28, EOG8F4VSG, EOG80306V, EOG8BCH22, EOG8ZKN28, EOG8F4VSG). While olfactory receptor gene families have rapidly contracted (EOG8Q5C4Z, EOG8RZ1DX), subfamilies of odorant binding proteins and gustatory receptors have grown (see Sensory Ecology section below). The expansion of gustatory receptor subfamilies are associated with bitter receptors, likely reflecting host plant detection of nightshades (Solanaceae). Some gene families associated with plant detoxification and insecticide resistance have rapidly expanded (Glutathione S-transferases: EOG85TG3K, EOG85F05D, EOG8BCH22; UDP-glycosyltransferase: EOG8BCH22; cuticle proteins: EOG8QJV4S, EOG8DNHJQ; ABC transporters: EOG83N9TJ), whereas others have contracted (Cytochrome P450s: EOG83N9TJ). Finally, gene families associated with immune defense (fibrinogen: EOG8DNHJQ; Immunoglobulin: EOG8Q87B6, EOG8S7N55, EOG854CDX, EOG8KSRZK, EOG8CNT5M) exhibit expansions that may be linked to defense against pathogens and parasitoids that commonly attack exposed herbivores [61]. A substantial proportion of the rapidly evolving gene families include proteins with transposable element domains (25.3%), while other important functional groups include DNA and protein binding (including many transcription factors), nuclease activity, protein processing, and cellular transport.

Diversification of transcription factor (TF) families potentially signals greater complexity of gene regulation, including enhanced cell specificity and refined spatiotemporal signaling [62]. Notably, several TF families are substantially expanded in *L. decemlineata*, including HTH_psq (194 genes vs. a mean of only 24 across the insects shown in **Figure 2**), MADF (152 vs. 54), and THAP (65 vs. 41). Two of these TF families, HTH_psq and THAP, are DNA binding domains in transposons. Of the 1,237 *L. decemlineata* TFs, we could infer DNA binding motifs for 189 (15%) (**Supplementary Table 4S**), mostly based on DNA binding specificity data from *Drosophila melanogaster* (124 TFs), but also from species as distant as human (45 TFs) and mouse (11 TFs). Motifs were inferred for a substantial proportion of the TFs in the largest TF families, including Homeodomain (59 of 90, 66%), bHLH (34 of 46, 74%), and Forkhead box (14 of 16, 88%). We could only infer a small number of C2H2 zinc finger motifs (21 of 439, ~5%), which is expected as these sequences evolve quickly by shuffling zinc finger arrays, resulting in largely dissimilar DNA-binding domains across metazoans [63]. Collectively, the almost 200 inferred DNA binding motifs for *L. decemlineata* TFs provide a unique resource to begin unraveling gene regulatory networks in this organism.

To identify genes active in mid-gut tissues, life-stages, and sex differences, we examined differential transcript expression levels using RNA sequencing data. Comparison of significantly differentially expressed genes with >100-fold change, after Bonferroni correction, indicated higher expression of digestive enzymes (proteases, peptidases, dehydrogenases and transporters) in mid-gut versus whole larval tissues, while cuticular proteins were largely expressed at lower levels (**Figure 3A, Supplementary Table 5S**). Comparison of an adult male and female showed higher expression of testes and sperm related genes in males, while genes involved in egg production and sterol biosynthesis are more highly expressed in females (**Figure 3B, Supplementary Table 6S**). Comparisons of larvae to both an adult male (**Figure 3C, Supplementary Table 7S**) and an adult female (**Figure 3D, Supplementary Table 8S**) showed higher expression of larval-specific cuticle proteins, and lower expression of odorant binding and chemosensory proteins. The adults, both drawn from a pesticide resistant population, showed higher constitutive expression of cytochrome p450 genes compared to the larval population, which is consistent with the results from previous studies of neonicotinoid resistance in this population [48].

**Figure 3.**
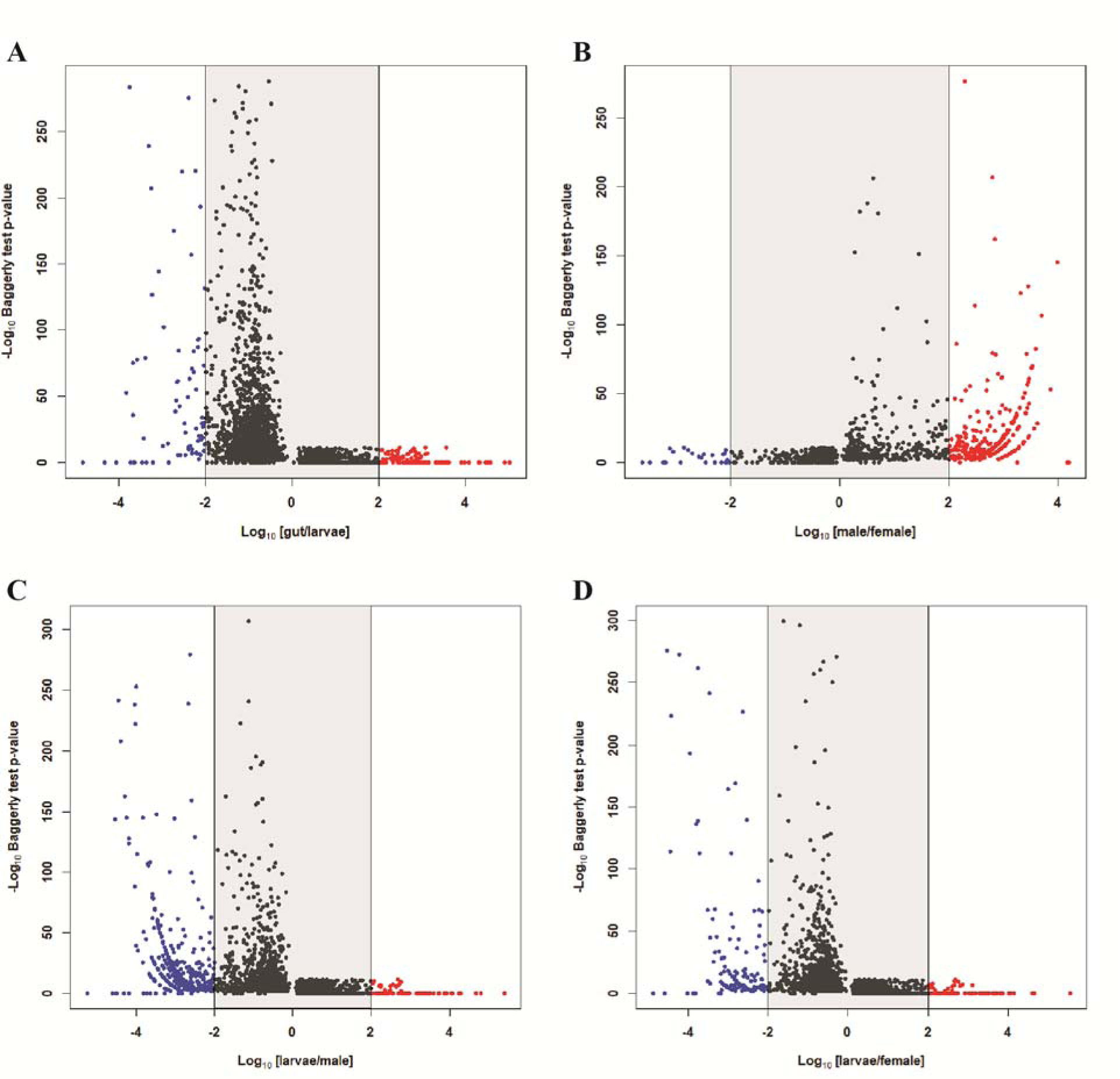
Volcano plots showing statistically significant gene expression differences after Bonferroni correction in *Leptinotarsa decemlineata*. A) Mid -gut tissue versus whole larvae, B) an adult male versus an adult female, C) an adult male versus whole larvae, and D) an adult female versus whole larvae. Points outside the gray area indicate >100-fold-differences in expression. Blue points indicate down-regulated genes and red points indicate up-regulated genes in each contrast.

### Transposable Elements

Transposable elements (TEs) are ubiquitous mobile elements within most eukaryotic genomes and play critical roles in both genome architecture and the generation of genetic variation [64]. Through insertional mutagenesis and recombination, TEs are a major contributor to the generation of novel mutations (50-80% of all mutation events within the genome of *D. melanogaster*) [65–67], and are increasingly thought to generate much of the genetic diversity that contributes to rapid evolution [68–70]. In addition to finding that genes with TE domains comprise 25% of the rapidly evolving gene families within *L. decemlineata*, we found that at least 17% of the genome consists of TEs (**Supplementary Table 9S**). This is substantially greater than the 6% found in *T. castaneum* [51], but less than some Lepidoptera (35% in *Bombyx mori* and 25% in *Heliconius melpomene*) [71,72]. LINEs were the largest TE class, comprising ~10% of the genome, while SINEs were not detected. Curation of the TE models with intact protein domains resulted in 334 current models of potentially active TEs, meaning that these TEs are capable of transposition and excision. Within the group of active TEs, we found 191 LINEs, 99 DNA elements, 38 LTRs, and 5 Helitrons. Given that TEs have been associated with the ability of species to rapidly adapt to novel selection pressures [73–75], particularly via alterations of gene expression patterns in neighboring genomic regions, we scanned gene rich regions (1 kb neighborhood size) for active TE elements. Genes with active neighboring TEs have functions that include transport, protein digestion, diapause, and metabolic detoxification (**Supplementary Table 10S**). Because TE elements have been implicated in conferring insecticide resistance in other insects [76], future work should investigate the role of these TE insertions on rapid evolutionary changes within pest populations of *L. decemlineata*.

### Population Genetic Variation and Invasion History

To understand the propensity for *L. decemlineata* pest populations to rapidly evolve across a range of environmental conditions, we examined geographical patterns of genomic variability and the evolutionary history of *L. decemlineata* (to see the genomic analysis of environmental adaptation, refer to the Diapause and Environmental Stress section of **Supplementary File**). High levels of standing variation are one mechanism for rapid evolutionary change [77]. We identified 1.34 million biallelic single nucleotide polymorphisms (SNPs) from pooled RNAseq datasets, or roughly 1 variable site for every 22 base pairs of coding DNA. This rate of polymorphism is exceptionally high when compared to vertebrates (e.g. ~1 per kb in humans, or ~1 per 500 bp in chickens) [78,79], and is 8-fold higher than other beetles (1 in 168 for *D. ponderosae* and 1 in 176 bp for *O. taurus)* [52,80] and 2 to 5-fold higher than some dipterans (1 in 54 bp for *D. melanogaster* and 1 in 125 bp for *Anopheles gambiae)* [78,81]. It is likely that these values simply scale with effective population size, although the dipterans, with the largest known population sizes, have reduced variation due to widespread selective sweeps and genetic bottlenecks [82].

Evolutionary relationships and the amount of genetic drift among Midwestern USA, Northeastern USA, and European *L. decemlineata* populations were estimated based on genome-wide allele frequency differences using a population graph. A substantial amount of local genetic structure and high genetic drift is evident among all populations, although both the reference lab strain from New Jersey and European populations appear to have undergone more substantial drift, suggestive of strong inbreeding (**Figure 4**). Population genetic divergence (*F_ST_*) values (**Supplementary Table 11S**) range from 0.035 (among Wisconsin populations) to 0.182 (Wisconsin compared to Europe). The allele frequency spectrum was calculated for populations in Wisconsin, Michigan, and Europe to estimate the population genetic parameter *θ*, or the product of the mutation rate and the ancestral effective population size, and the ratio of contemporary to ancestral population size in models that allowed for single or multiple episodes of population size change. Estimates of ancestral *θ* are much higher for Wisconsin (*θ* = 12595) and Michigan (*θ* = 93956) than Europe (*θ* = 3.07; **Supplementary Table 12S**), providing support for a single introduction into Europe following a large genetic bottleneck [15]. In all three populations, a model of population size growth is supported, in agreement with historical accounts of the beetles expanding from the Great Plains into the Midwestern U.S. and Europe [3,15], but the dynamics of each population appear independent, with the population from Michigan apparently undergoing a very recent decline in contemporary population size (the ratio of contemporary to ancient population size is 0.066, compared to 3.3 and 2.1 in Wisconsin and Europe, respectively).

**Figure 4.**
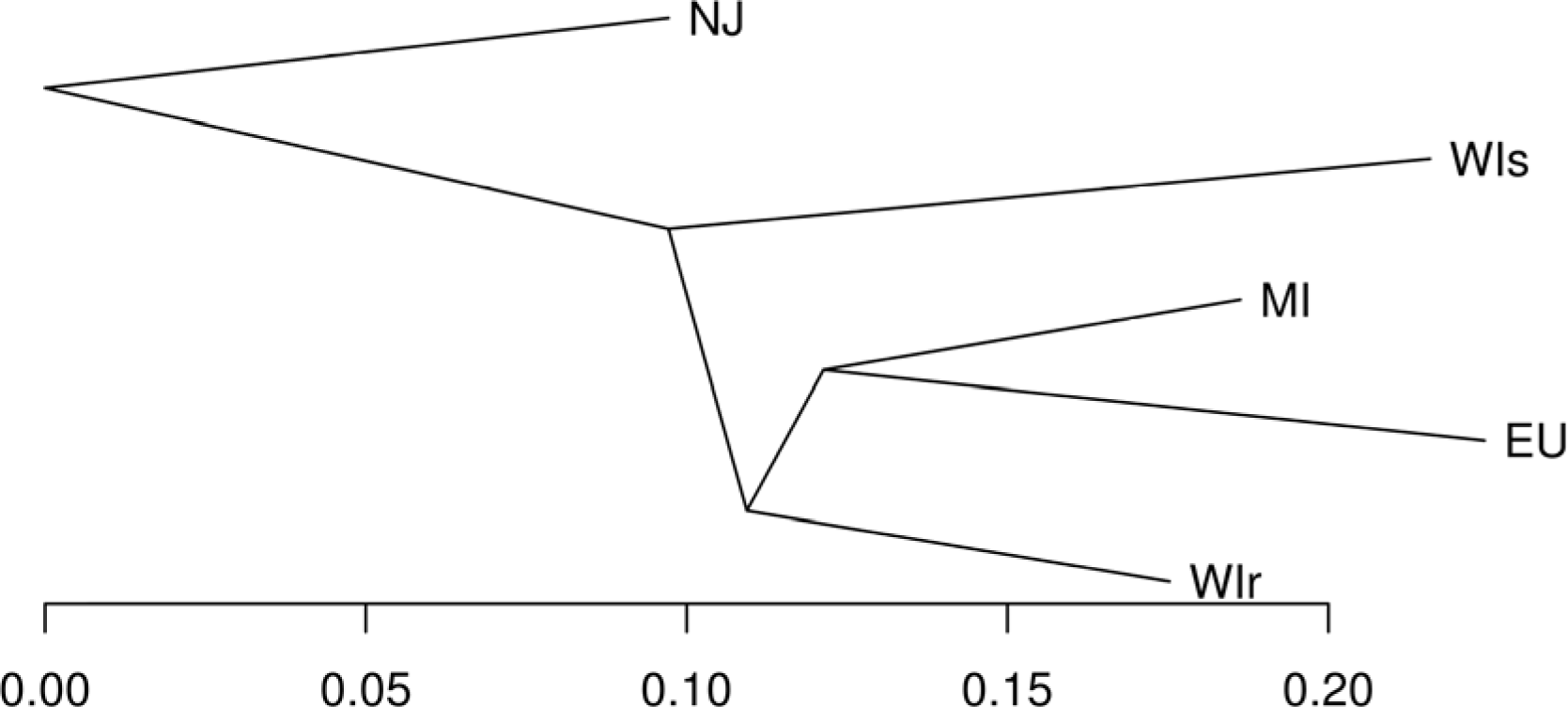
Population genetic relationships and relative rates of genetic drift among *Leptinotarsa decemlineata* pest populations based on single nucleotide polymorphism data. Population codes: NJ-New Jersey lab strain, WIs-imidacloprid susceptible population from Arlington, Wisconsin, WIr-imidacloprid resistant population from Hancock, Wisconsin, MI-imidacloprid resistant population from Michigan, and EU-European samples combined from Italy and Russia.

### Sensory Ecology and Host Plant Detection

To interact with their environment, insects have evolved neurosensory organs to detect environmental signals, including tactile, auditory, chemical and visual cues [83]. We examined neural receptors, olfactory genes, and light sensory (opsin) genes to understand the sensory ecology and host-plant specializations of *L. decemlineata*.

We found high sequence similarity in the neuroreceptors of *L. decemlineata* compared to other Coleoptera. The transient receptor potential (TRP) channels are permeable transmembrane proteins that respond to temperature, touch, pain, osmolarity, pheromones, taste, hearing, smell and visual cues of insects [84,85]. In most insect genomes, there are typically 13-14 TRP genes located in insect stretch receptor cells and several are targeted by commercial insecticides [86]. We found 12 TRP genes present in the *L. decemlineata* genome, including the two TRPs (Nanchung and Inactive) that are targeted by commercial insecticides, representing a complete set of one-to-one orthologs with *T. castaneum*. Similarly, the 20 known amine neurotransmitter receptors in *T. castaneum* are present as one-to-one orthologs in *L. decemlineata* [87,88]. Amine receptors are G-protein-coupled receptors that interact with biogenic amines, such as octopamine, dopamine and serotonin. These neuroactive substances regulate behavioral and physiological traits in animals by acting as neurotransmitters, neuromodulators and neurohormones in the central and peripheral nervous systems [89].

The majority of phytophagous insects are restricted to feeding on several plant species within a genus, or at least restricted to a particular plant family [90]. Thus, to find their host plants within heterogeneous landscapes, insect herbivores detect volatile organic compounds through olfaction, which utilizes several families of chemosensory gene families, such as the odorant binding proteins (OBPs), odorant receptors (ORs), gustatory receptors (GRs), and ionotropic receptors (IRs) [91]. OBPs directly bind with volatile organic compounds emitted from host plants and transport the ligands across the sensillar lymph to activate the membrane-bound ORs in the dendrites of the olfactory sensory neurons [92]. The ORs and GRs are 7-transmembrane proteins related within a superfamily [93] of ligand-gated ion channels [94]. The ionotropic receptors are related to ionotropic glutamate receptors and function in both smell and taste [95]. These four gene families are commonly large in insect genomes, consisting of tens to hundreds of members.

We compared the number of genes found in *L. decemlineata* in the four chemosensory gene families to *T. castaneum* and *A. glabripennis* (Table 2, as well as **Supplementary Tables 13S-16S** for details). While the OBP family is slightly enlarged, the three receptor families are considerably smaller in *L. decemlineata* than in either *A. glabripennis* or *T. castaneum,* consistent with the specialization of this beetle on one genus of plants. However, each beetle species exhibits species-specific gene subfamily expansions (**Supplementary Figures 5S-8S**); in particular, some lineages of GRs related to bitter taste are expanded in *L. decemlineata* relative to *A. glabripennis* and other beetles. Among the OBPs, we identified a major *L. decemlineata*-specific expansion of proteins belonging to the Minus-C class (OBPs that have lost two of their six conserved cysteine residues) that appear unrelated to the ‘traditional’ Minus-C subfamily in Coleoptera, indicating that coleopteran OBPs have lost cysteines on at least two occasions.

**Table 2.** Numbers of putatively functional proteins in four chemosensory families in *Leptinotarsa decemlineata* and other beetle species.

| Species | Odorant Binding Proteins (OBPs) | Olfactory Receptors (ORs) | Gustatory Receptors (GRs) | Ionotropic Receptors (IRs) |
| --- | --- | --- | --- | --- |
| *Leptinotarsa decemlineata* | 58 | 75 | 144 | 27 |
| *Anoplophora glabripennis* | 52 | 120 | 234 | 72 |
| *Tribolium castaneum* | 49 | 264 | 219 | 72 |

To understand the visual acuity of *L. decemlineata*, we examined the G-protein-coupled transmembrane receptor opsin gene family. We found five opsins, three of which are members of rhabdomeric opsin (R-opsin) subfamilies expressed in the retina of insects [51,96]. Specifically, the *L. decemlineata* genome contains one member of the long wavelength-sensitive R-opsin and two short wavelength UV-sensitive R-opsins. The latter were found to be closely linked in a range of less than 20,000 bp, suggestive of recent tandem gene duplication. Overall, the recovered repertoire of retinally-expressed opsins in *L. decemlineata* [97] is consistent with the beetle’s attraction to yellow light and to the yellow flowers of its ancestral host plant, *S. rostratum* [98,99], and is consistent with the beetle’s sensitivity in the UV- and LW-range [100]. In addition, we found a member of the Rh7 R-opsin subfamily, which is broadly conserved in insects including other beetle species (*A. glabripennis*), although it is missing from *T. castaneum*. Finally, *L. decemlineata* has a single ortholog of the c-opsin subfamily shared with *T castaneum*, which is absent in *A. glabripennis* and has an unclear role in photoreception [101].

### Host Plant Utilization

#### Protein digestion

Insect herbivores are fundamentally limited by nitrogen availability [102], and thus need to efficiently break down plant proteins in order to survive and develop on host plants [103]. *Leptinotarsa decemlineata* has serine and cysteine digestive peptidases (coined “intestains”) [32,104,105], as well as aspartic and metallo peptidases [33], for protein digestion. For the vast majority of plant-eating beetles (the infraorder Cucujiformia, which includes Chrysomelidae), cysteine peptidases contribute most strongly to proteolytic activity in the gut [105,106]. In response to herbivory, plants produce a wide range of proteinase inhibitors to prevent insect herbivores from digesting plant proteins [103,105,107]. Coleopteran peptidases are differentially susceptible to plant peptidase inhibitors, and our annotation results suggest that gene duplication and selection for inhibitor insensitive genotypes may have contributed to the success of leaf-feeding beetles (Chrysomelidae) on different plants. We found that gene expansion of cysteine cathepsins from the C1 family in *L. decemlineata* correlates with the acquisition of greater digestive function by this group of peptidases, which is supported by gene expression activity of these genes in mid-gut tissue (**Figure 3A, Figure 5**). The gene expansion may be explained by an evolutionary arms race between insects and plants that favors insects with a variety of digestive peptidases in order to resist plant peptidase inhibitors [108,109] and allows for functional specialization [110].

**Figure 5.**
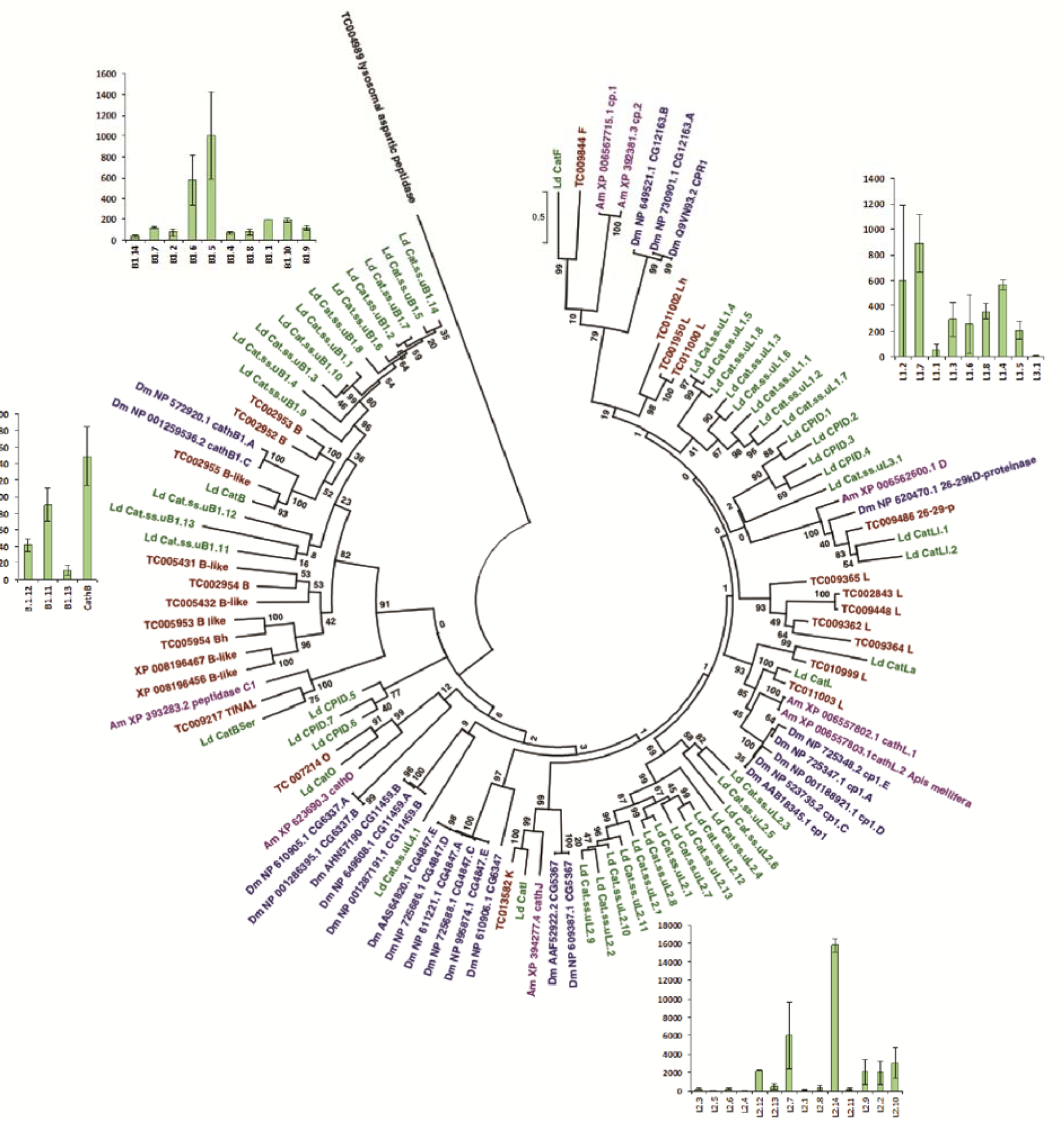
Phylogenetic relationships of the cysteine peptidase gene family in *Leptinotarsa decemlineata* compared to model insects. Species abbreviations are: *L. decemlineata* (Ld, green color), *Drosophila melanogaster* (Dm, blue color), *Apis mellifera* (Am, purple color), and *Tribolium castaneum* (Tc, red color). Mid-gut gene expression (TPM) of highly expressed *L. decemlineata* cysteine peptidases is shown as bar graphs across three replicate treatments.

Cysteine peptidases of the C1 family were represented by more than 50 genes separated into four groups with different structure and functional characteristics (**Supplementary Table 18S**): cathepsin L subfamily, cathepsin B subfamily, TINAL-like genes, and cysteine peptidase inhibitor domains (CPIDs). Cathepsin L subfamily cysteine peptidases are endopeptidases [111] that can be distinguished by the cathepsin propeptide inhibitor domain I29 (pfam08246) [112,113]. Within the cathepsin L subfamily, we found sequences that were similar to classical cathepsin L, cathepsin F, and cathepsin O. However, there were 28 additional predicted peptidases of this subfamily that could not be assigned to any of the “classical” cathepsin types, and most of these were grouped into two gene expansions (uL1 and uL2) according to their phylogenetic and structural characteristics. Cathepsin B subfamily cysteine peptidases are distinguished by the specific peptidase family C1 propeptide domain (pfam08127). Within the cathepsin B subfamily, there was one gene corresponding to typical cathepsin B peptidases and 14 cathepsin B-like genes. According to the structure of the occluding loop, only the typical cathepsin B may have typical endo- and exopeptidase activities, while a large proportion of cathepsin B-like peptidases presumably possesses only endopeptidase activity due to the absence of a His-His active subsite in the occluding loop, which is responsible for exopeptidase activity [111]. Only one gene corresponding to a TINAL-like-protein was present, which has a domain similar to cathepsin B in the C-terminus, but lacks peptidase activity due to the replacement of the active site Cys residue with Ser [114]. Cysteine peptidase inhibitor domain (CPID) genes encode the I29 domain of cysteine peptidases without the mature peptidase domain. Within the CPID group, there were seven short inhibitor genes that lack the enzymatic portion of the protein. A similar trend of “stand-alone inhibitors” has been observed in other insects, such as *B. mori* [115]. These CPID genes may be involved in the regulation of cysteine peptidases. We note that we found multiple fragments of cysteine peptidase genes, suggesting that the current list of *L. decemlineata* genes may be incomplete. Comparison of these findings with previous data on *L. decemlineata* cysteine peptidases [116] demonstrates that intestains correspond to several peptidase genes from the uL1 and uL2 groups (**Supplementary Table 18S**). These data, as well as literature for Tenebrionidae beetles [117], suggest that intensive gene expansion is typical for peptidases that are involved in digestion.

We also found a high number of digestion-related serine peptidase genes in the *L. decemlineata* genome (**Supplementary Table 19S**), but they contribute only a small proportion of the beetle’s total gut proteolytic activity [32]. Of the 31 identified serine peptidase genes and fragments, we annotated 16 as trypsin-like peptidases and 15 as chymotrypsin-like peptidases. For four chymotrypsin-like and one trypsin-like peptidase, we identified only short fragments. All complete (and near-complete) sequences have distinctive S1A peptidase subfamily motifs, a conserved catalytic triad, conserved sequence residues such as the "CWC" sequence and cysteines that form disulfide bonds in the chymotrypsin protease fold. The number of serine peptidases was higher than expected based upon the number of previously identified EST clones [32], but lower than the number of chymotrypsin and trypsin genes in the *T. castaneum* genome.

#### Carbohydrate digestion

Carbohydrates are the other category of essential nutrients for *L. decemlineata.* The enzymes that assemble and degrade oligo- and polysaccharides, collectively termed Carbohydrate active enzymes (CAZy), are categorized into five major classes: glycoside hydrolases (GH), polysaccharide lyases, carbohydrate esterases (CE), glycosyltransferases (GT) and various auxiliary oxidative enzymes [118]. Due to the many different roles of carbohydrates, the CAZy family profile of an organism can provide insight into “glycobiological potential” and, in particular, mechanisms of carbon acquisition [119]. We identified 182 GHs assigned to 25 families, 181 GTs assigned to 41 families, and two CEs assigned to two families in *L. decemlineata*; additionally, 99 carbohydrate-binding modules (which are non-catalytic modules associated with the above enzyme classes) were present and assigned to 9 families (**Supplementary Table 20S**; the list of CAZy genes is presented in **Supplementary Table 21S**). We found that *L. decemlineata* has three families of genes associated with plant cell wall carbohydrate digestion (GH28, GH45 and GH48) that commonly contain enzymes that target pectin (GH28) and cellulose (GH45 and GH48), the major structural components of leaves [120]. We found evidence of massive gene duplications in the GH28 family (14 genes) and GH45 family (11 genes, plus one additional splicing variant), whereas GH48 is represented by only three genes in the genome [121]. Overall, the genome of *L. decemlineata* shows a CAZy profile adapted to metabolize pectin and cellulose contained in leaf cell walls. The absence of specific members of the families GH43 (α-L-arabinofuranosidases) and GH78 (α-L-rhamnosidases) suggests that *L. decemlineata* can break down homogalacturonan, but not substituted galacturonans such as rhamnogalacturonan I or II [120]. The acquisition of these plant cell wall degrading enzymes has been linked to horizontal transfer in the leaf beetles and other phytophagous beetles [120], with strong phylogenetic evidence supporting the transfer of GH28 genes from a fungal donor (Pezizomycotina) in *L. decemlineata,* as well as in the beetles *D. ponderosae* and *Hypothenemus hampei,* but a novel fungal donor in the more closely related cerambycid beetle *A. glabripennis* [54] and a bacterial donor in the weevil *Callosobruchus maculatus* [120].

### Insecticide Resistance

To understand the functional genomic properties of insecticide resistance, we examined genes important to neuromuscular target site sensitivity, tissue penetration, and prominent gene families involved in Phase I, II, and III metabolic detoxification of xenobiotics [122]. These include the cation-gated nicotinic acetylcholine receptors (nAChRs), the γ-amino butyric acid (GABA)-gated anion channels and the histamine-gated chloride channels (HisCls), cuticular proteins, cytochrome P450 monooxygenases (CYPs), and the Glutathione S-transferases (GSTs).

Many of the major classes of insecticides (organochlorides, organophosphates, carbamates, pyrethroids, neonicotinoids and ryanoids) disrupt the nervous system (particularly ion channel signaling), causing paralysis and death [123]. Resistance to insecticides can come from point mutations that reduce the affinity of insecticidal toxins to ligand-gated ion superfamily genes [124]. The cys-loop ligand-gated ion channel gene superfamily is comprised of receptors involved in mediating synaptic ion flow during neurotransmission [125]. A total of 22 cys-loop ligand-gated ion channels were identified in the *L. decemlineata* genome in numbers similar to those observed in other insects [126], including 12 nAChRs, three GABA receptors, and two HisCls (**Supplementary Figure 9S**). The GABA-gated chloride channel homolog of the *Resistance to dieldrin* (Rdl) gene of *D. melanogaster* was examined due to its role in resistance to dieldrin and other cyclodienes in Diptera [127]. The coding sequence is organized into 10 exons (compared to nine in *D. melanogaster*) on a single scaffold, with duplications of the third and sixth exon (**Supplementary Figure 10S**). Alternative splicing of these two exons encodes for four different polypeptides in *D. melanogaster* [128,129], and as the splice junctions are present in *L. decemlineata,* we expect the same diversity of *Rdl*. The point mutations in the transmembrane regions TM2 and TM3 of *Rdl* are known to cause insecticide resistance in Diptera [124,127], but were not observed in *L. decemlineata*.

Cuticle genes have been implicated in imidacloprid resistant *L. decemlineata* [130] and at least one has been shown to have phenotypic effects on resistance traits following RNAi knockdown [131]. A total of 163 putative cuticle protein genes were identified and assigned to one of seven families (CPR, CPAP1, CPAP3, CPF, CPCFC, CPLCG, and TWDL) (**Supplemental Figure 11S** and **Table 22S**). Similar to other insects, the CPR family, with the RR-1 (soft cuticle), RR-2 (hard cuticle), and unclassifiable types, constituted the largest group of cuticle protein genes (132) in the *L. decemlineata* genome. While the number of genes in *L. decemlineata* is slightly higher than in *T. castaneum* (110), it is similar to *D. melanogaster* (137) [132]. Numbers in the CPAP1, CPAP3, CPF, and TWDL families were similar to other insects, and notably no genes with the conserved sequences for CPLCA were detected in *L. decemlineata,* although they are found in other Coleoptera.

A total of 89 CYP (P450) genes were identified in the *L. decemlineata* genome, an overall decrease relative to *T. castaneum* (143 genes). Due to their role in insecticide resistance in *L. decemlineata* and other insects [38,48,133], we examined the CYP6 and CYP12 families in particular. Relative to *T. castaneum*, we observed reductions in the CYP6BQ, CYP4BN, and CYP4Q subfamilies. However, five new subfamilies (CYP6BJ, CYP6BU, CYP6E, CYP6F and CYP6K) were identified in *L. decemlineata* that were absent in *T. castaneum*, and the CYP12 family contains three genes as opposed to one gene in *T. castaneum* (CYP12h1). We found several additional CYP genes not present in *T. castaneum*, including CYP413A1, CYP421A1, CYP4V2, CYP12J and CYP12J4. Genes in CYP4, CYP6, and CYP9 are known to be involved in detoxification of plant allelochemicals as well as resistance to pesticides through their constitutive overexpression and/or inducible expression in imidacloprid resistant *L. decemlineata* [48,130].

GSTs have been implicated in resistance to organophosphate, organochlorine and pyrethroid insecticides [134] and are responsive to insecticide treatments in *L. decemlineata* [130,135]. A total of 27 GSTs were present in the *L. decemlineata* genome, and while they represent an expansion relative to *A. glabripennis,* all have corresponding homologs in *T. castaneum*. The cytosolic GSTs include the epsilon (11 genes), delta (5 genes), omega (4 genes), theta (2 genes) and sigma (3 genes) families, while two GSTs are microsomal (**Supplementary Figure 12S**). Several GST-like genes present in the *L. decemlineata* genome represent the Z class previously identified using transcriptome data [135].

### Pest Control via the RNAi Pathway

RNA interference (RNAi) is the process by which small non-coding RNAs trigger sequence-specific gene silencing [136], and is important in protecting against viruses and mobile genetic elements, as well as regulating gene expression during cellular development [137]. The application of exogenous double stranded RNA (dsRNA) has been exploited as a tool to suppress gene expression for functional genetic studies [138,139] and for pest control [41,140].

We annotated a total of 49 genes associated with RNA interference, most of them were found on a single scaffold. All genes from the core RNAi machinery (from all three major RNAi classes) were present in *L. decemlineata*, including fifteen genes encoding components of the RNA Induced Silencing Complex (RISC) and genes known to be involved in double-stranded RNA uptake, transport, and degradation (**Supplementary Table 22S**). A complete gene model was annotated for *R2D2*, an essential component of the siRNA pathway that interacts with *dicer-2* to load siRNAs into the RISC, and not previously detected in the transcriptome of the *L. decemlineata* mid-gut [141]. The core components of the small interfering RNA (siRNA) pathway were duplicated, including *dicer-2*, an RNase III enzyme that cleaves dsRNAs and pre-miRNAs into siRNAs and miRNAs respectively [142,143]. The *dicer-2a* and *dicer-2b* CDS have 60% nucleotide identity to each other, and 56% and 54% identity to the *T. castaneum dicer-2* homolog, respectively. The *argonaute-2* gene, which plays a key role in RISC by binding small non-coding RNAs, was also duplicated. A detailed analysis of these genes will be necessary to determine if the duplications provide functional redundancy. The duplication of genes in the siRNA pathway may play a role in the high sensitivity of *L. decemlineata* to RNAi knockdown [144] and could benefit future efforts to develop RNAi as a pest management technology.

### Conclusion

The whole-genome sequence of *L. decemlineata*, provides novel insights into one of the most diverse animal taxa, Chrysomelidae. It is amongst the largest beetle genomes sequenced to date, with a minimum assembly size of 640 Mb (ranging up to 1.17 Gb) and 24,740 genes. The genome size is driven in part by a large number of transposable element families, which comprise at least 17% of the genome and appear to be rapidly expanding relative to other beetles. Population genetic analyses suggest high levels of nucleotide diversity, local geographic structure, and evidence of recent population growth, which helps to explain how *L. decemlineata* rapidly evolves to exploit novel host plants, climate space, and overcome a range of pest management practices (including a large and diverse number of insecticides). Digestive enzymes, in particular the cysteine peptidases and carbohydrate-active enzymes, show evidence of gene expansion and elevated expression in gut tissues, suggesting the diversity of the genes is a key trait in the beetle’s phytophagous lifestyle. Additionally, expansions of the gustatory receptor subfamily for bitter tasting might be a key adaptation to exploiting hosts in the nightshade family, Solanaceae, while expansions of novel subfamilies of CYP and GST proteins are consistent with rapid, lineage-specific turnover of genes implicated in *L. decemlineata’s* capacity for insecticide resistance. Finally, *L. decemlineata* has interesting duplications in RNAi genes that might increase its sensitivity to RNAi and provide a promising new avenue for pesticide development. The *L. decemlineata* genome promises new opportunities to investigate the ecology, evolution, and management of this species, and to leverage genomic technologies in developing sustainable methods of pest control.

## Materials and Methods

### Genome Characteristics and Sampling of DNA and RNA

Previous cytological work determined that *L. decemlineata* is diploid and consists of 34 autosomes plus an XO system in males, or an XX system in females [145]. Twelve chromosomes are submetacentric, while three are acrocentric and two are metacentric, although one chromosome is heteromorphic (acrocentric and/or metacentric) in pest populations [146]. The genome size has been estimated with Feulgen densitometry at 0.46 pg, or approximately 460 Mb [47]. To generate a reference genome sequence, DNA was obtained from a single adult female, sampled from an imidacloprid resistant strain developed from insects collected from a potato field in Long Island, NY. Additionally, whole-body RNA was extracted for one male and one female from the same imidacloprid resistant strain. Raw RNAseq reads for 8 different populations were obtained from previous experiments: two Wisconsin populations (PRJNA297027) [130], a Michigan population (PRJNA400685), a lab strain originating from a New Jersey field population (PRJNA275431), and three samples from European populations (PRJNA79581 and PRJNA236637) [43,121]. All RNAseq data came from pooled populations or were combined into a population sample from individual reads. In addition, RNA samples of an adult male and female from the same New Jersey population were sequenced separately using Illumina HiSeq 2000 as 100 bp paired end reads (deposited in the GenBank/EMBL/DDBJ database, PRJNA275662), and three samples from the mid-gut of 4^th^-instar larvae were sequenced using SOLiD 5500 Genetic Analyzer as 50 bp single end reads (PRJNA400633).

### Genome Sequencing, Assembly, Annotation and Assessment

Four Illumina sequencing libraries were prepared, with insert sizes of 180 bp, 500 bp, 3 kb, and 8 kb, and sequenced with 100 bp paired-end reads on the Illumina HiSeq 2000 platform at estimated 40x coverage, except for the 8kb library, which was sequenced at estimated 20x coverage. ALLPATHS-LG v35218 [49] was used to assemble reads. Two approaches were used to scaffold contigs and close gaps in the genome assembly. The reference genome used in downstream analyses was generated with ATLAS-LINK v1.0 and ATLAS GAP-FILL v2.2 (https://www.hgsc.bcm.edu/software/). In the second approach, REDUNDANS was used [57], as it is optimized to deal with heterozygous samples. The raw sequence data and *L. decemlineata* genome have been deposited in the GenBank/EMBL/DDBJ database (Bioproject accession PRJNA171749, Genome assembly GCA_000500325.1, Sequence Read Archive accessions: SRX396450-SRX396453). This data can be visualized, along with gene models and supporting data, at the i5k Workspace@NAL: https://i5k.nal.usda.gov/Leptinotarsa_decemlineata and https://apollo.nal.usda.gov/lepdec/jbrowse/ [147]. We estimated the size of our genome using a kmer distribution plot in JELLYFISH [148], where we mapped the 100 bp paired-end reads from the 180 bp insert library, used a 19 bp kmer distribution plot, and corrected for ploidy.

Automated gene prediction and annotation were performed using MAKER v2.0 [149], using RNAseq evidence and arthropod protein databases. The MAKER predictions formed the basis of the first official gene set (OGS v0.5.3). To improve the structural and functional annotation of genes, these gene predictions were manually and collaboratively edited using the interactive curation software Apollo [150]. For a given gene family, known insect genes were obtained from model species, especially *T. castaneum* [51] and *D. melanogaster* [151], and the nucleotide or amino acid sequences were used in *blastx* or *tblastn* [152,153] to search the *L. decemlineata* OGS v0.5.3 or genome assembly, respectively, on the i5k Workspace@NAL. All available evidence (AUGUSTUS, SNAP, RNA data, etc.), including additional RNAseq data not used in the MAKER predictions, were used to inspect and modify gene predictions. Changes were tracked to ensure quality control. Gene models were inspected for quality, incorrect splits and merges, internal stop codons, and gff3 formatting errors, and finally merged with the MAKER-predicted gene set to produce the official gene set (OGS v1.0; merging scripts are available upon request). For focal gene families (e.g. peptidase genes, odorant and gustatory receptors, RNAi genes, etc.), details on how genes were identified and assigned names based on functional predictions or evolutionary relationship to known reference genes are provided in the Supplementary Material (**Supplementary File**).

To assess the quality of our genome assemblies, we used BUSCO v2.0 [59] to determine the completeness of each genome assembly and the official gene set (OGS v1.0), separately. We benchmarked our data against 35 insect species in the Endopterygota obd9 database, which consists of 2,442 single-copy orthologs (BUSCOs). Secondly, we annotated and examined the genomic architecture of the Hox and Iroquois Complex gene clusters. For this, tBLASTn searches were performed against the genome using orthologous Hox gene protein sequences from *T. castaneum* (Tcas3.0) and *A. glabripennis.* Provisional *L. decemlineata* models were refined, and potential gene duplications were identified, via iterative and reciprocal BLAST and by manual inspection and correction of protein alignments generated with ClustalW2 [154], using RNAseq expression evidence when available. Finally, we used BlobTools v1.0 [155] to assess the assembled genome for possible contamination by generating a Blobplot. This plot integrates guanine and cytosine (GC) content of sequences, read coverage and sequence similarity via blast searches to assess genome contamination. Putative contaminants were identified using DIAMOND BLAST [156] to query the genome scaffolds against the UniProt reference database [157], and read coverage was assessed by mapping the 100 bp paired-end reads from the 180 bp insert library to the ALLPATHS genome with the Burrows-Wheeler Aligner (bwa-mem) version 0.7.5a [158].

### Gene Family Evolution

In order to identify rapidly evolving gene families along the *L. decemlineata* lineage, we obtained ~38,000 ortho-groups from 72 Arthropod species as part of the i5k pilot project (Thomas et al. *in prep)* from OrthoDB version 8 [159]. For each ortho-group, we took only those genes present in the order Coleoptera, which was represented by the following six species: *A. glabripennis, A. planipennis, D. ponderosae, L. decemlineata, O. taurus,* and *T. castaneum* (http://i5k.github.io/) [51,52,54]. Finally, in order to make accurate inferences of ancestral states, families that were present in only one of the six species were removed. This resulted in a final count of 11,598 gene families that, among these six species, form the comparative framework that allowed us to examine rapidly evolving gene families in the *L. decemlineata* lineage.

Aside from the gene family count data, an ultrametric phylogeny is also required to estimate gene gain and loss rates. To make the tree, we considered only gene families that were single copy in all six species and that had another arthropod species also represented with a single copy as an outgroup. Outgroup species were ranked based on the number of families in which they were also single copy along with the coleopteran species, and the highest ranking outgroup available was chosen for each family. For instance, *Pediculus humanus* was the most common outgroup species. For any gene family, we chose *P. humanus* as the outgroup if it was also single copy. If it was not, we chose the next highest ranking species as the outgroup for that family. This process resulted in 3,932 single copy orthologs that we subsequently aligned with PASTA [160]. We used RAxML [161] with the PROTGAMMAJTTF model to make gene trees from the alignments and ASTRAL [162] to make the species tree. ASTRAL does not give branch lengths on its trees, a necessity for gene family analysis, so the species tree was again given to RAxML along with a concatenated alignment of all one-to-one orthologs for branch length estimation. Finally, to generate an ultrametric species tree with branch lengths in millions of years (my) we used the software r8s [163], with a calibration range based on age estimates of a crown Coleopteran fossil at 208.5-411 my [164]. This calibration point itself was estimated in a similar fashion in a larger phylogenetic analysis of all 72 Arthropod species (Thomas et al. *in prep*).

With the gene family data and ultrametric phylogeny as the input data, gene gain and loss rates (*Λ*) were estimated with CAFE v3.0 [165]. CAFE is able to estimate the amount of assembly and annotation error (*ε*) present in the input data using a distribution across the observed gene family counts and a pseudo-likelihood search, and then is able to correct for this error and obtain a more accurate estimate of *λ.* Our analysis had an *ε* value of about 0.02, which implies that 3% of gene families have observed counts that are not equal to their true counts. After correcting for this error rate, *λ* = 0.0010 is on par with those previously those found for other Arthropod orders (Thomas et al. *in prep*). Using the estimated *λ* value, CAFE infers ancestral gene counts and calculates p-values across the tree for each family to assess the significance of any gene family changes along a given branch. Those branches with low p-values are considered rapidly evolving.

### Gene Expression Analysis

RNAseq analyses were conducted to 1) establish male, female, and larva-enriched gene sets and 2) identify specific genes that are enriched within the digestive tract (mid-gut) compared to entire larva. RNAseq datasets were trimmed with CLC Genomics v.9 (Qiagen) and quality was assessed with FastQC (http://www.bioinformatics.babraham.ac.uk/projects/fastqc). Each dataset was mapped to the predicted gene set (OGS v1.0) using CLC Genomics. Reads were mapped with >90% similarity over 60% of length, with two mismatches allowed. The number of reads were corrected to reads per million mapped to allow comparison between RNAseq datasets that having varying coverage. Transcripts per million (TPM) was used as a proxy for gene expression and fold changes were determined as the TPM in one sample relative to the TPM of another dataset [166]. The Baggerly’s test (t-type test statistic) followed by Bonferroni correction was used to identify genes with significant enrichment in a specific sample [167]. Statistical values for Bonferroni correction were reported as the number of genes x *α* value. This stringent statistical analysis was used as only a single replicate was available for each treatment. Enriched genes were removed, and mapping and expression analyses were repeated to ensure low expressed genes were not missed. Genes were identified by BLASTx searching against the NCBI non-redundant protein databases for arthropods with an expectation value (E-value) < 0.001.

### Transposable Elements

We investigated the identities (family membership), diversity, and genomic distribution of active transposable elements within *L. decemlineata* in order to understand their contribution to genome structure and to determine their potential positional effect on genes of interest (focusing on a TE neighborhood size of 1 kb). To identify TEs and analyze their distribution within the genome, we developed three repeat databases using: 1) RepeatMasker [168], which uses the library repeats within Repbase (http://www.girinst.org/repbase/), 2) the program RepeatModeler [169], which identifies de-novo repeat elements, and 3) literature searches to identify beetle transposons that were not found within Repbase. The three databases were used within RepeatMasker to determine the overall TE content in the genome.

To eliminate false positives and examine the genome neighborhood surrounding active TEs, all TE candidate models were translated in 6 frames and scanned for protein domains from the Pfam and CDD database (using the software transeq from Emboss, hmmer3 and rps-blast). The protein domain annotations were manually curated in order to remove: a) clear false positives, b) old highly degraded copies of TEs without identifiable coding potential, and c) the correct annotation when improper labels were given. The TE models that contained protein domains were mapped onto the genome and used for our neighborhood analysis: we extracted the 1 kb flanking regions for each gene and scanned these regions for TEs with intact protein coding domains.

### Population Genetic Variation and Demographic Analysis

Population genetic diversity of pooled RNAseq samples was used to examine genetic structure of pest populations and past population demography. For Wisconsin, Michigan and the lab strains from New Jersey, we aligned the RNAseq data to the genomic scaffolds, using Bowtie2 version 2.1.0 [170] to index the genome and generate aligned SAM files. We used bwa [150] to align the RNAseq from the three populations from Europe. SAMtools/BCFtools version 0.1.19 [171] was used to produce BAM and VCF files. All calls were filtered with VCFtools version 0.1.11 [172] using a minimum quality score of 30 and minimum depth of 10. All indels were removed from this study. Population specific VCF files were sorted and merged using VCFtools, and the allele counts were extracted for each SNP. These allele frequency data were then used to infer population splits and relative rates of genetic drift using Treemix version 1. 12 [173]. We ran Treemix with SNPs in groups of 1000, choosing to root the tree with the Wisconsin population. In addition, for each pair of populations, we estimated the average genetic divergence by using F-statistics (*F_ST_*) in VCFtools to calculate the ratio of among- to within-population genetic divergence across SNP loci.

To infer patterns of demographic change in the Midwestern USA (Wisconsin and Michigan) and European populations, the genome-wide allele frequency spectrum was used in *dadi* version 1.6.3 [174] to infer demographic parameters under several alternative models of population history. The history of *L. decemlineata* as a pest is relatively well-documented. The introduction of *L. decemlineata* into Europe in 1914 [175] is thought to have involved a strong bottleneck [15] followed by rapid expansion. Similarly, an outbreak of *L. decemlineata* in Nebraska in 1859 is thought to have preceded population expansion into the Midwest reaching Wisconsin in 1865 [3]. For each population, a constant-size model, a two-epoch model of instantaneous population size change at a time point τ, a bottle-growth model of instantaneous size change followed by exponential growth, and a three-epoch model with a population size change of fixed duration followed by exponential growth, was fit to infer *Θ,* the product of the ancestral effective population size and mutation rate, and relative population size changes.

## Acknowledgements

We sincerely thank the sequencing, assembly and annotation teams at the Baylor College of Medicine Human Genome Sequencing Center for their efforts. We would like to acknowledge the following funding sources: sequencing, assembly and automated annotation was supported by NIH grant NHGRI U54 HG003273 to RAG; the UVM Agricultural Experiment Station Hatch grant to YHC (VT-H02010); the NIH postdoctoral training grant to RFM (K12 GM000708); MMT’s work with Apollo was supported by NIH grants (5R01GM080203 from NIGMS, and 5R01HG004483 from NHGRI) and by the Director, Office of Science, Office of Basic Energy Sciences, of the U.S. Department of Energy (contract No. DE-AC02-05CH11231); the National Science Centre (2012/07/D/NZ2/04286) and Ministry of Science and Higher Education scholarship to AM. Mention of trade names or commercial products in this publication is solely for the purpose of providing specific information and does not imply recommendation or endorsement by the U.S. Department of Agriculture. USDA is an equal opportunity provider and employer.

## Authors’ contributions

MNA, AB, JHB, KC, AKC, OC, EMD, ENE, PE, MF, IG-R, CG, AG, MG, BH, ECJ, JWJ, MK, SAK, AK, FL, VL, XM, AM, NJM, RFM, BO, SRP, KAP, YP, LCP, MP, ÉR, JPR, HMR, AJR, VMR-A, GS, AST, IMVJ, ADY, GDY, and J-SY contributed to the manual annotation effort, data analysis, and data interpretation, in addition to reading and approving the final manuscript. SDS and YHC coordinated the project and drafted the manuscript. SR and RAG coordinated genome sequencing, assembly and automated annotation at the Baylor College of Medicine Human Genome Sequencing Center. MFP, CC, MMT, and M-JMC coordinated the biocuration of the genome. GWCT generated the phylogeny and conducted the gene family analysis. MTW conducted the transcription factor analysis. AG, KG, MP, and SZ contributed additional RNAseq data. JBB coordinated the RNAseq data analysis. KB, AM, and YHC conducted the transposable element analysis. SDS and JC conducted the population genetics analysis.

## Competing interests

The authors declare that they have no competing interests.

## Availability of data and material

All data generated or analyzed during this study have been made publicly available (see Methods for NCBI accession numbers), or included in this published article and its supplementary information. The genome assembly and official gene sets can be accessed at: https://data.nal.usda.gov/dataset/leptinotarsa-decemlineata-genome-assembly-10_5667, https://data.nal.usda.gov/dataset/leptinotarsa-decemlineata-genome-annotations-v053_5668 and https://data.nal.usda.gov/dataset/leptinotarsa-decemlineata-official-gene-set-v11.

## Supplementary Files

Supplementary File.pdf Supplementary Materials. Contains additional methods, results, figures and tables.

Supplementary Dataset 1. Precursor miRNA nucleotide sequences.

Supplementary Dataset 2. miRNA predictions (gff format).

Supplementary Dataset 3. Peptide sequences of annotated olfactory genes. Includes the Odorant Binding Proteins (OBPs), Odorant Receptors (ORs), Gustatory Receptors (GRs), and Ionotropic Receptors (IRs). The IRs from *Tribolium castaneum* are also included.

